# Achilles tendons of MRL/MpJ mice show scar-mediated healing after tenotomy

**DOI:** 10.1101/2025.05.13.653102

**Authors:** Ningfeng Tang, Divya S. Sivalingam, Kimberly Wilson, Mehari M. Weldemariam, Hongying Tian, Christoph D. Hart, Prishasai Ramnath, Jacob Glazier, Jie Jiang, Natalie Leong, Tao Lowe, Maureen A. Kane, Masahiro Iwamoto, Catherine K. Kuo, Motomi Enomoto-Iwamoto

**Author notes:** Corresponding Authors: Catherine K. Kuo, Fischell Department of Bioengineering, A. James Clark School of Engineering, University of Maryland, 4108 A. James Clark Hall, 8278 Paint Branch Dr, College Park, MD 20742, USA, Motomi Enomoto-Iwamoto, Department of Orthopaedics, University of Maryland School of Medicine, 670 W Baltimore St, HSFIII 7171, Baltimore, MD 21201, USA.

## Abstract

Tendon healing in Murphy Ross Large (MRL/MpJ) mice was examined using an Achilles tendon tenotomy model, a full transection model, compared to a patella focal injury model in which tendons in MRL/MpJ mice showed regenerative healing. After complete transection, the Achilles tendons of MRL/MpJ mice healed via scar formation, regardless of sex, as did C57BL/6J mice. Tensile testing found that mechanical properties of injured tendons of both MRL/MpJ female and male mice exhibited comparable levels to those of C57BL/6J female and male mice, respectively after 4 weeks of healing, which is during the remodeling phase. After 10 weeks of healing, injured tendons of MRL/MpJ mice possessed smaller heterotopic ossification volumes than those of C57BL/6J mice. Proteomics analysis revealed similar alterations to signaling pathways in injured tendons of MRL/MpJ and C57BL/6J male mice after 4 weeks of healing. However, among the altered pathways, actin cytoskeleton and integrin signaling pathways were two of the top pathways that were more prominently activated in C57BL/6J males than in MRL/MpJ males. These findings indicate that MRL/MpJ mice possess limited capacity to regenerate injured tendons after complete rupture, and that tenotomized Achilles tendons of MRL/MpJ male mice have lesser induction of heterotopic ossification and lower activation of the signaling pathways also induced with injury in C57BL/6J male mice.

## 1. Introduction

Tendons have limited repair capacity. Injured tendons do not usually regain their original levels of biological, structural, and mechanical characteristics, and instead can undergo chronic degeneration ^1–5^. In contrast to embryonic and neonatal tendons ^6,7^, adult tendons heal via scar formation characterized by high cell density, disorganized cell arrangement, thin and randomly aligned collagen fibers, high vascularity, and mucoid matrix ^5,8^. Scar formation inhibits recovery of mechanical properties. Failure of full functional recovery may result in chronic tendon disorders and re-rupture, leading to poor quality of life and work disability ^5^. Murphy Roths Large (MRL/MpJ) mice have strong repair capacity in musculoskeletal tissues including articular cartilage, muscle, and bone ^9^. MRL/MpJ mice also exhibit more regenerative healing of focal injuries of the patellar tendon in adults, based on better recovery of mechanical properties and improved extracellular matrix and cell alignment when compared to healing of the same injury in C57BL/6 mice ^10,11^. A flexor tendon injury model involving partial laceration has also shown that MRL/MpJ tendons heal with reduced fibrotic adhesion ^12^.

Based on these compelling prior works, we were interested in investigating whether tendons in MRL/MpJ mice undergo regenerative healing even in an Achilles tendon complete transection model, which is different from the focal injury models previously used to study tendon repair in MRL/MpJ mice ^10–12^. Determining the molecular mechanism that supports the reparative ability of the MRL/MpJ is crucial to develop new modalities to promote effective tendon healing, and thus we also used proteomics analyses to examine which signaling pathways may be involved in improved tendon repair by MRL/MpJ mice.

## 2. Materials and Methods

### 2.1. Mouse experiments

All animal studies were conducted with approval from the Institutional Animal Care and Use Committee at the University of Maryland, Baltimore (0120007 and 1122006). Animals were housed in a University Laboratory Animal Resources supervised animal facility with a 12-hour light/dark cycle and controlled temperature of 22 ± 1°C and humidity of 55 ± 5%. Animals were provided hygienic animal bedding of wood shavings and cotton pads. The health status of each animal was monitored throughout the experiments by investigators, animal veterinary technicians, and veterinarians, as per institutional guidelines. The mice were free of all viral, bacterial, and parasitic pathogens during the experimental schedule.

A complete transverse transection (tenotomy), without attempt at repair, was made at the midpoint of the left Achilles tendon in 12-week-old female and male C57BL/6J (#000646, The Jackson Laboratory, Bar Harbor, ME) and MRL/MpJ (#000486, The Jackson Laboratory) mice under anesthesia. The analgesic carprofen, 4.5 mg/Kg was administrated pre- and post-surgery. The mice receiving the tenotomy surgery were euthanized 4 or 10-weeks post-surgery, which was during the early and late remodeling phase, respectively. The right and left hind limbs were collected for histological analysis and mechanical tests. The mid-substance of Achilles tendons was snap frozen in liquid nitrogen and stored at -80°C until use. The criteria for animal exclusion during the experiment, determined prior to the experiment, were to exclude mice that received analgesics and/or antibiotics and/or were euthanized due to infection or distress. No mice were excluded in this study. No exclusion criteria were set for data points during the analysis. No data points were excluded. MRL/MpJ and C57BL/6J mice were maintained in the same location and compared, but randomization was not used and confounders were not controlled. A total of 109 mice were used: 3 female C57BL/6J, 3 male C57BL/6J, 3 female MRL/MpJ, and 3 male MRL/MpJ mice for histological examination at 4 weeks post-surgery; 8 female C57BL/6J, 5 female MRL/MpJ, 8 male C57BL/6J, and 7 female MRL/MpJ mice for mechanical testing at 4 weeks post-surgery; 5 male C57BL/6J, 5 male MRL/MpJ, 4 female C57BL/6J, and 3 female MRL/MpJ mice for μCT analysis at 10 weeks post-surgery; 18 uninjured male C57BL/6J mice (for 5 unique biological replicates), 10 injured male C57BL/6J mice (for 5 unique biological replicates), 10 uninjured male MRL/MpJ mice (for 4 unique biological replicates), and 8 injured male MRL/MpJ mice (for 4 unique biological replicates) for proteomics at 4-weeks post-surgery; and 3 C57BL/6J and 3 MRL/MpJ male mice for immunoblotting at 4 weeks post-surgery. Healing ability comparisons between MRL/MpJ and C57BL/6J mice were based on mechanical properties (primary outcome measure) and histology (secondary outcome measure).

### 2.2. Histology

The hind limbs were fixed with 4% paraformaldehyde for 24 hours at 4°C and decalcified with 10% ethylenediaminetetraacetic acid at pH 7.2. Serial sagittal sections (5 μm) of the injured or contralateral uninjured Achilles tendons were made. Sections were stained with hematoxylin-eosin and picrosirius red. Hematoxylin-eosin staining images were captured using light microscopy (BZX-700, Keyence, Tokyo, Japan). Picrosirius red staining images were captured using polarized light microscopy (20X objective, ZEISS Axio Scan, Germany) ^7,13–15^.

### 2.3. Micro-computed tomography (microCT) analysis

Samples were scanned using a Skyscan 1172 microCT scanner (Bruker, Billerica, MA, USA) while immersed in phosphate buffered saline without Ca^2+^ or Mg^2+^ (PBS). Specifically, images were acquired using previously reported conditions ^16^: 6.5 µm/pixel; 0.2 mm Al filter using a 55 kV X-ray source; resolution at 2 K with step rotation of 0.51; and two average frames using 360° rotation. Image reconstruction was performed using NRecon software (Bruker). We manually identified regions of heterotopic ossification by setting threshold between 80 and 255, for which volume analyses were performed using CTAn software (Bruker). Rendering images were obtained using CTVOX (Bruker).

### 2.4. Mass spectrometry-based proteomics

We collected tendon samples and stored them at -80 °C until assay. Lysis buffer containing 5% SDS (sodium dodecyl sulfate) and 50 mM triethylammonium bicarbonate (1 M, pH 8.0) was added to tendons and the samples were then homogenized using bead beating with a Precellys CK14 lysing kit (Bertin Corp., Rockville, MD). Next, protein extraction and digestion were performed as described ^17^ using S-trap micro columns (ProtiFi, NY). Digested peptides were quantified using a BCA assay kit (Thermo Fisher Scientific, 23275) to determine peptide concentration. LC-MS/MS-based proteomic analysis was conducted on a nanoACQUITY Ultra-Performance Liquid Chromatography system (Waters Corporation, Milford, MA USA) coupled to an Orbitrap Fusion Lumos Tribrid mass spectrometer (Thermo Scientific, San Jose, CA USA). Peptide separation was effected on a nanoACQUITY Ultra-Performance Liquid Chromatography (UPLC) analytical column (BEH130 C18, 1.7 µm, 75 µm x 200 mm; Waters Corporation, Milford, MA, USA) using a 185-min linear gradient with 3-40% acetonitrile and 0.1% formic acid. Mass spectrometry conditions were as follows: Full MS scan resolution of 240,000, precursor ions fragmentation by high-energy collisional dissociation of 35%, and a maximum cycle time of 3 seconds. The resulting mass spectra were processed using Thermo Proteome Discoverer ((PD, version 3.0.0.757), Thermo Fisher Scientific) and searched against a UniProt mouse reference proteome (release 2024.03, 17180 entries) using Sequest HT algorithm. Search parameters include carbamidomethylation of cysteines (+57.021 Da) as a static modification, methionine oxidation (+15.995 Da) as a dynamic modification, precursor mass tolerance of 20 ppm, fragment mass tolerance of 0.5 Da, and trypsin as a digestion enzyme. Tryptic missed cleavages were restricted to a maximum of two, with peptide lengths set between 6 and 144 residues. For protein quantification, the Minora feature detector, integrated in the PD, was used as described previously ^17^. To ensure high data quality, proteins were further filtered to a 1% false discovery rate (FDR) threshold, calculated with the Percolator algorithm. Next, protein abundance values exported from PD were post-processed using Perseus software (version 1.6.14.0). Proteins with missing values were excluded to improve data quality. Differentially expressed proteins (DEPs) were identified using a two-tailed Student’s t-test (adjusted p-value < 0.05). Ingenuity Pathway Analysis (IPA) software (Qiagen, Germantown, MD) was used to identify dysregulated pathways and biological processes. The data were deposited in the ProteomeXchange via the PRIDE database, with the project accession number PXD057712.

### 2.6. Mechanical Testing

Achilles tendons with calcanei were fine dissected to ensure removal of all musculature and surrounding soft tissue. Injured and contralateral uninjured tendons were hydrated in PBS immediately after dissection and throughout the course of mechanical testing. Prior to mechanical testing, the dissected tendons were suspended in a tube pinching the calcaneus with the lid and scanned using a Skyscan 1172 microCT scanner (Camera pixel size, 9.00; Rotation range, 360 deg; Image rotation, 3 deg; scanning time, about 9 mins). PBS was placed in the bottom of the tube to prevent the tendon from drying. The volume and length of the tendon were measured using CTAn software. The cross-sectional area (CSA) was then calculated as the ratio of volume to length. Mechanical testing was performed, as previously described ^18^, on a DMA850 dynamic mechanical analyzer (TA Instruments, New Castle, DE) with TRIOS Software. Tendons were gripped between the calcaneus and myotendinous junction and underwent a uniaxial tensile testing protocol consisting of a preload to 0.02 N, and then ramp to failure test at 0.1% strain per second. Mechanical (ultimate force, stiffness) and material (elastic modulus, ultimate stress, ultimate strain) properties were determined, as previously described ^19^.

### 2.7. Statistical analysis

Power analysis (G*Power v.3.1, Heinrich Heine Universität Düsseldorf, Germany) performed using experimental data, with power of 0.80 and significance level α=0.05, determined that a minimum sample size of N=5 biological replicates (different mice) per experimental group was needed. Data was assumed to be drawn from a normally distributed population. Normality was analyzed by the Shapiro-Wilk test. Non-parametric tests were selected when the assumption of normality could not be met. Mechanical testing and heterotopic ossification results were analyzed using Prism 10 (10.6.0, Graphpad, San Diego, CA). Material and mechanical properties were compared between sexes, mouse strains, and uninjured and injured tendons using a one-way ANOVA with Tukey’s post-hoc. Heterotopic ossification results were analyzed by Mann-Whitney or Kologorov-Smirnov test to identify differences between groups. The threshold for significance for all tests was set as p<0.05. The data met the assumptions of the statistical approach.

## 3. Results

### 3.1. Both C57BL/6J and MRL/MpJ mouse Achilles tendons presented with scar formation after tenotomy

C57BL/6J and MRL/MpJ mice received unilateral tenotomy of the Achilles tendon. To investigate an early remodeling phase, we examined the recovery of extracellular matrix organization at 4 weeks after surgery. Sagittal sections of injured Achilles tendons from both female and male mice were acquired. Figure 1 shows sagittal sections of polarized images stained with picrosirius red from the distal calcaneal insertion (left, single arrow) to the proximal myotendinous junction (right, double arrow). Injured C57BL/6J tendons showed disorganized immature collagen fibers (Fig. 1A and 1E), compared to aligned collagen fibers in uninjured tendons of age- and sex-matched mice (Fig. 1B and 1F). Injured Achilles tendons of both female and male MRL/MpJ mice also showed disorganized immature collagen fibers with polarized imaging (Fig. 1C and 1G), in contrast to uninjured tendons (Fig. 1D and 1H). The hematoxylin and eosin-stained sections of injured tendons presented as scar tissue rich in fibroblasts and blood vessels (Fig. 2A–2H), whereas uninjured Achilles tendons from age-, sex-, and strain-matched mice appeared dense with extracellular matrix and aligned tenocytes. (Fig. 2I–2L).

**Fig. 1.**
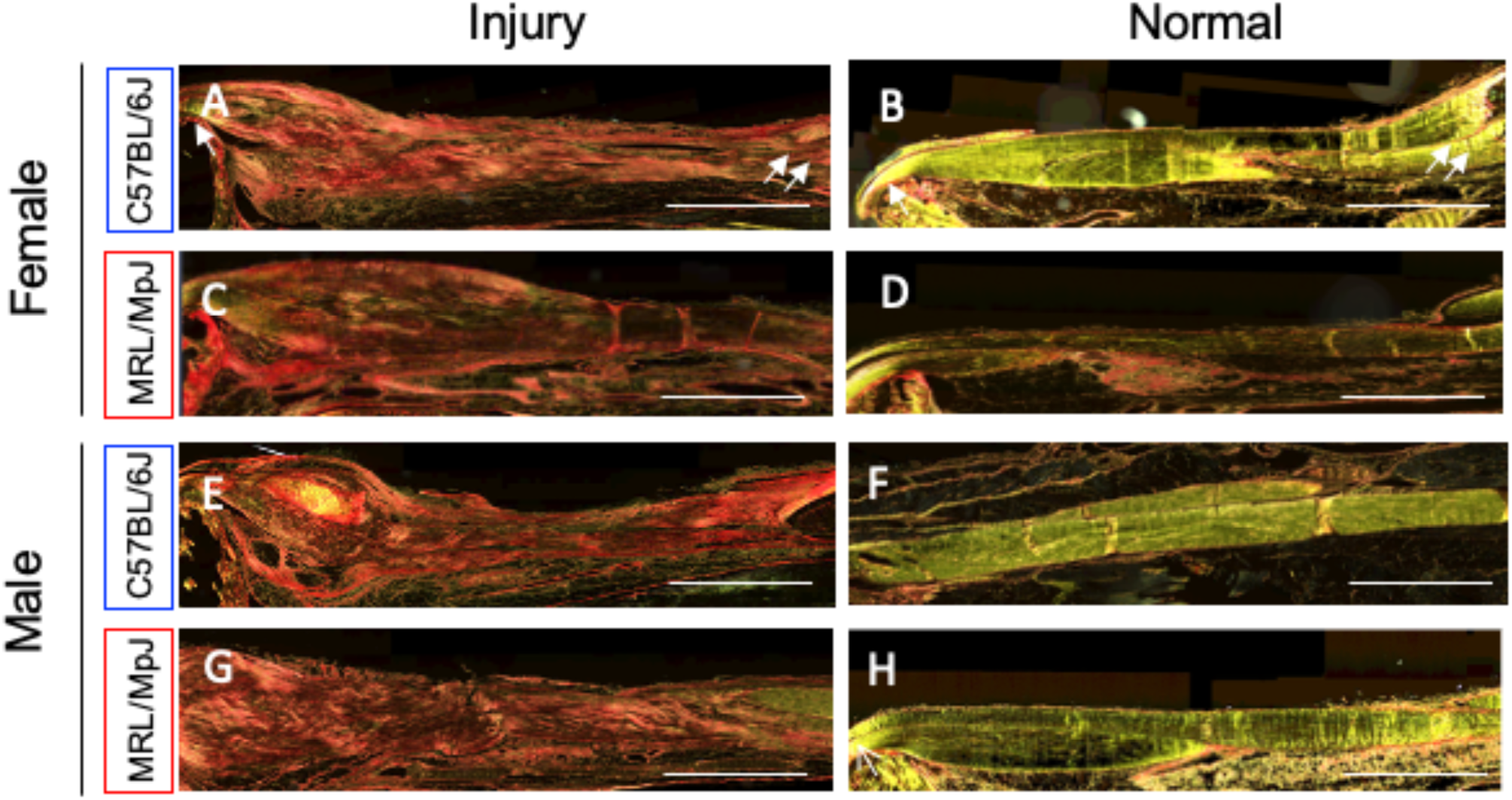
Comparison of picrosirius red (PSR)-stained sections of injured tendons between C57BL/6J and MRL/MpJ mice. A complete transverse transection was made at the midpoint of the left Achilles tendon in 12-week-old female and male C57BL/6J (A, B, E, and F) and MRL/MpJ (C, D, G, and H) mice (n=3 per sex and strain). Sagittal sections of the injured (A, C, E, and G) and contralateral uninjured (B, D, F, and H) Achilles tendons four weeks after surgery were stained with PSR. Bar represents 1.5 mm. Arrow and double arrow show distal calcaneal insertion and proximal myotendinous junction of the Achilles tendon, respectively.

**Fig. 2.**
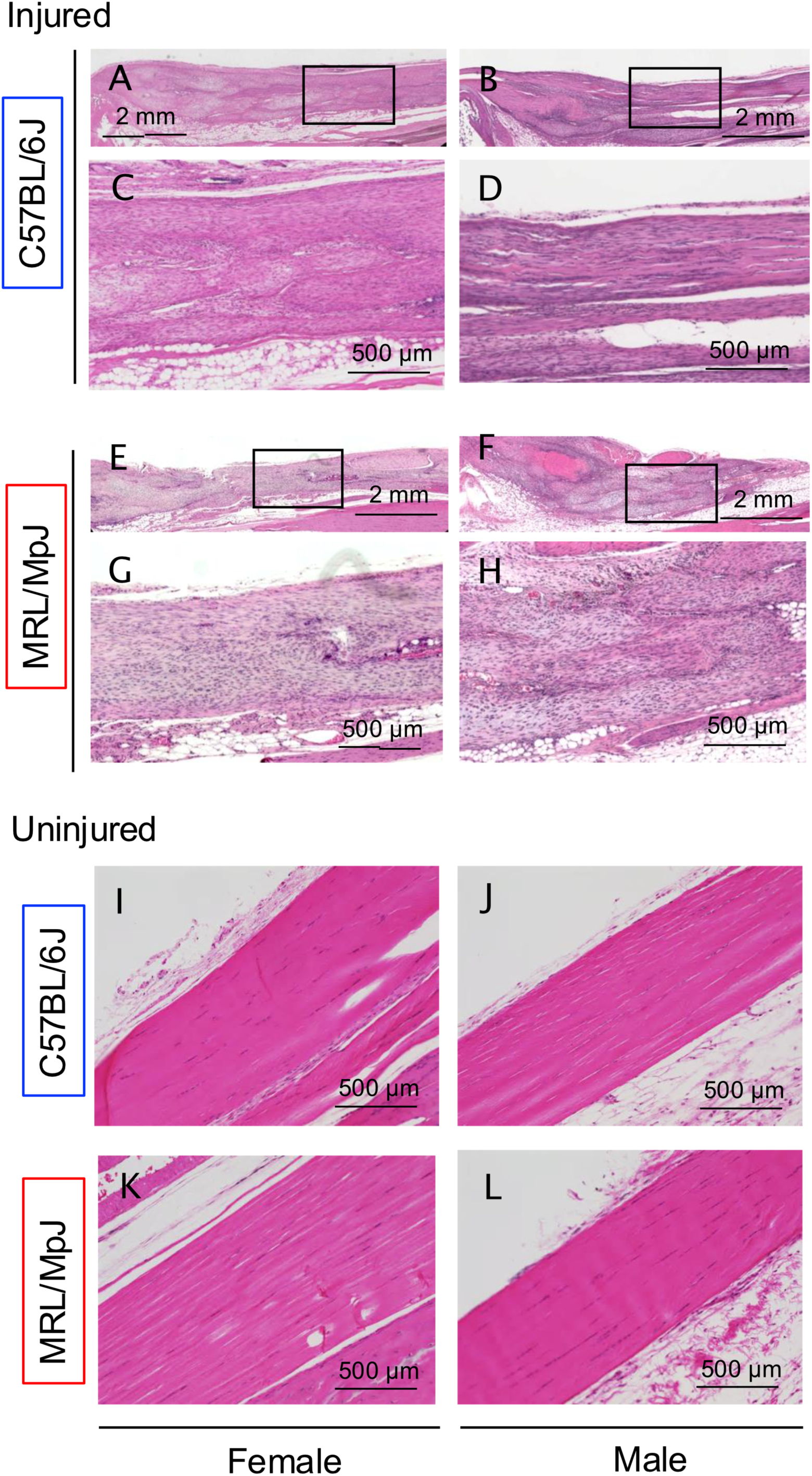
Comparison of hematoxylin-eosin-stained sections of injured tendons between C57BL/6J and MRL/MpJ mice. A complete transverse transection was made at the midpoint of the left Achilles tendon in 12-week-old female and male C57BL/6J (A-D, I and J) and MRL/MpJ (E-H, K and L) mice (n=3 per sex and strain). Sagittal sections of the injured (A-H) and contralateral uninjured (I-L) Achilles tendons four weeks after surgery were stained with hematoxylin-eosin (HE).

### 3.2. Recovery of material and mechanical properties of injured MRL/MpJ Achilles tendons were comparable to that of C57BL/6J Achilles tendons

Injured Achilles tendons of female and male MRL/MpJ mice possessed lower stiffness, elastic modulus, ultimate load, and ultimate stress, but greater ultimate strain compared to their corresponding uninjured MRL/MpJ Achilles tendons (Fig. 3, S1, and S2). Similarly, injured female and male C57BL/6J tendon stiffness, elastic modulus, ultimate load, and ultimate stress were lower, whereas ultimate strain was higher, than their corresponding uninjured Achilles tendons (Fig. 3 and S2). In addition, the injury-induced changes in stiffness, elastic modulus, ultimate load, ultimate stress, and ultimate strain were similar between MRL/MpJ and C57BL/6J tendons (Fig. 3 and S2), suggesting that recovery of material and mechanical properties of injured MRL/MpJ tendons is comparable to that of injured C57BL/6J tendons. Further, sex-specific differences in material and mechanical properties in either injured or uninjured tendons of either strain were not found (Fig. 3 and S2).

**Fig. 3.**
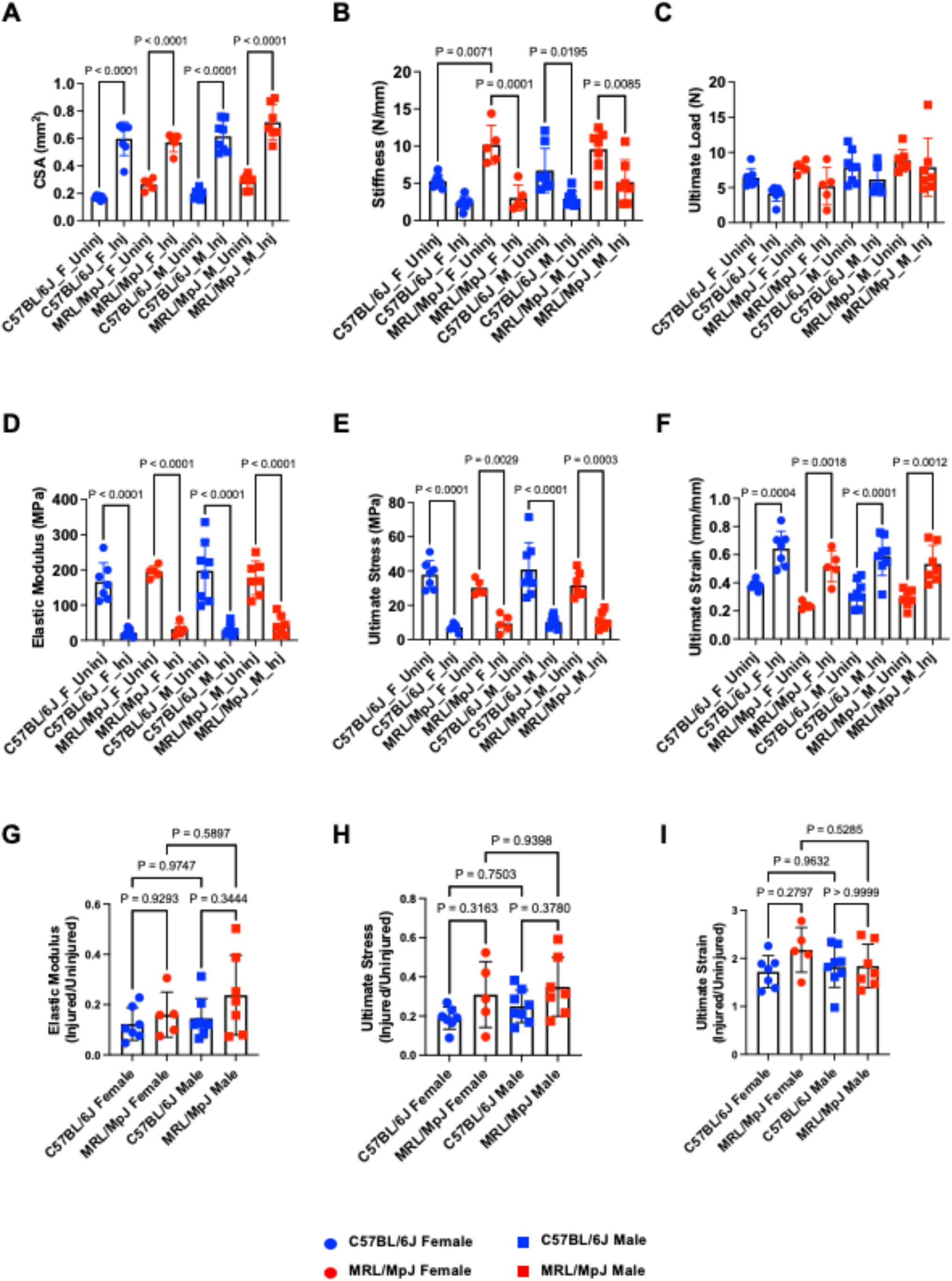
Comparison of biomechanical properties of injured tendons between C57BL/6J and MRL/MpJ mice. A complete transverse transection was made at the midpoint of the left Achilles tendon in 12-week-old female (circles; F) and male (squares; M) C57BL/6J (blue; n=7 for female and n=8 for male) and MRL/MpJ (red; n=5 for female and n=7 for male) mice. Four weeks after surgery, injured (Inj; left limb) and contralateral uninjured (Uninj; right limb) limbs were harvested. Dissected tendons were mechanically assessed using tensile testing. (A) Cross-sectional area (CSA), (B) stiffness, (C) ultimate load, (D) elastic modulus, (E) ultimate stress, and (F) ultimate strain were compared between uninjured and injured tendons of C57BL/6J female (blue circles), MRL/MpJ female (red circles), C57BL/6J male (blue squares), and MRL/MpJ male (red squares). Data points represent individual biological replicates. For (A-F), p-values of statistically significant comparisons between uninjured and injured of the same sex and strain, between sexes of the same injury status and strain, and between strains of the same injury status and sex are shown. Averages and standard deviations of absolute values are summarized in Fig. S2 of Supplementary Material. (G-I) Injured values were normalized to average uninjured values of the respective group and (G) elastic modulus, (H) ultimate stress, and (I) ultimate strain were compared between female and male C57BL/6J and MRL/MpJ mice. Data points represent individual biological replicates, with averages and standard deviations indicated. Normalized elastic modulus averages and standard deviations were 0.123 +/- 0.065 for C57BL/6J female, 0.160 +/- 0.090 for MRL/MpJ female, 0.145 +/- 0.079 for C57BL/6J male, and 0.238 +/- 0.159 for MRL/MpJ male. Normalized ultimate stress averages and standard deviations were 0.189 +/- 0.056 for C57BL/6J female, 0.310 +/- 0.168 for MRL/MpJ female, 0.250 +/- 0.083 for C57BL/6J male, and 0.349 +/- 0.149 for MRL/MpJ male. Normalized ultimate strain averages and standard deviations were 1.722 +/- 0.334 for C57BL/6J female, 2.177 +/- 0.466 for MRL/MpJ female, 1.826 +/- 0.431 for C57BL/6J male, and 1.838 +/- 0.455 for MRL/MpJ male. No statistically significant differences were found between normalized mechanical properties between strains of the same sex or sexes of the same strain.

### 3.3. MRL/MpJ Achilles tendons possessed lower volume of heterotopic ossification compared to C57BL/6J Achilles tendons

The mouse Achilles tendon tenotomy model has been studied as a heterotopic ossification model based on the established phenomenon of chondroid degeneration followed by ectopic bone formation in injured Achilles tendons by later timepoints of healing ^20^. Injury-induced heterotopic ossification was analyzed by microCT scan ten weeks after surgery, the time when heterotopic ossification is evident in this model ^20^. Injury induced heterotopic ossification in both C57BL/6J and MRL/MpJ Achilles tendons (Fig. 4A). The total volume of heterotopic ossified regions was smaller in MRL/MpJ male tendons than that in C57BL/6J male tendons (Fig. 4B, left). Combining male and female results also revealed significantly smaller volumes of heterotopic ossification in MRL/MpJ mice (Fig. 4B, right).

**Fig. 4.**
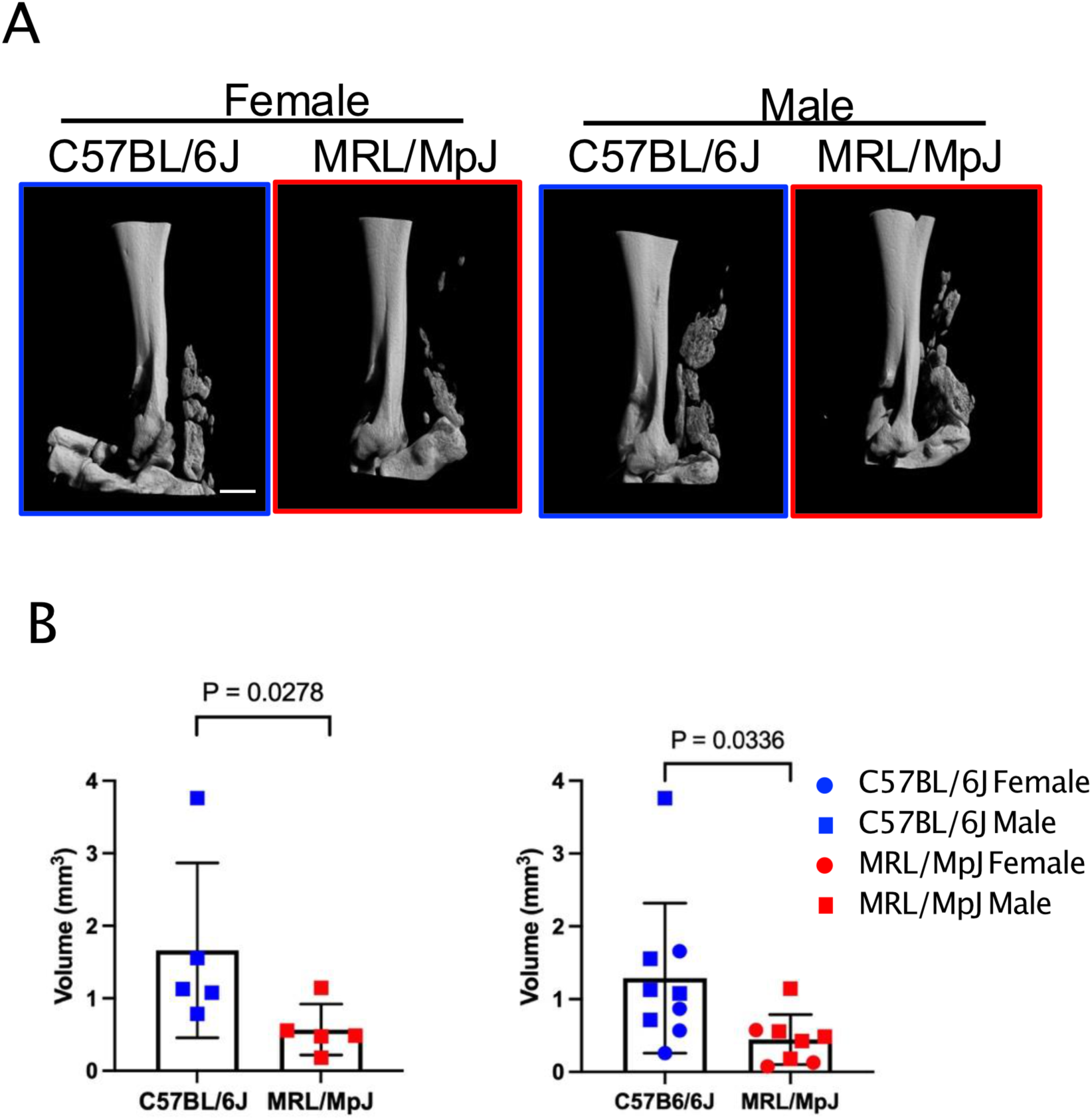
Comparison of heterotopic ossification in injured tendons between C57BL/6J and MRL/MpJ mice. A complete transverse transection was made at the midpoint of the left Achilles tendon in 12-week-old male (square) and female (circle) C57BL/6J (blue) and MRL/MpJ (red) mice (n=5 per strain). Ten weeks after surgery, injured (left) limbs were harvested. (A) Heterotopic ossification was imaged by microCT. Bar represents 1 mm. (B) Total volume of ossified regions within the injured tendon was analyzed. Data points represent individual biological replicates, with averages and standard deviations indicated. Averages and standard deviations were 1.65+/-1.219 for C57BL/6J male, 0.588+/-0.348 for MRL/MpJ male, 1.24+/- 0.573 for C57BL/6J male and female and 0.400+/-0.268 mm^3^ MRL/MpJ male and female mice.

### 3.4. Activation of actin cytoskeleton and integrin signaling pathways was less pronounced in MRL/MpJ male Achilles injured tendons

We performed proteomic analysis of the samples obtained from C57BL/6J and MRL/MpJ male injured and uninjured Achilles tendons. Differentially expressed proteins (DEPs) were identified using a two-tailed Student’s t-test with a criterion of adjusted p-value < 0.05 (Fig. 5A). We found that 538 proteins and 595 proteins were up-regulated in C57BL/6J injured tendons and MRL/MpJ injured tendons, respectively, compared with the mouse strain-matched uninjured tendons, whereas 79 proteins and 54 proteins were down-regulated in C57BL/6J injured tendons and MRL/MpJ injured tendons, respectively. Comparison between C57BL/6J and MRL/MpJ male injured tendons showed that 335 proteins were up-regulated and 70 proteins were down-regulated in C57BL/6J injured tendons compared to MRL/MpJ injured tendons. In spite of differences in DEPs between C57BL/6J and MRL/MpJ samples, based on IPA, C57BL/6J and MRL/MpJ injured tendons revealed similar alterations of the canonical pathways relative to their corresponding uninjured tendons (Supplementary Table 1). When the top 100 pathways were compared between C57BL/6J and MRL/MpJ groups, the top 50 pathways overlapped completely, and exhibited differences only in the rank order (common pathways in Supplementary Table 1, highlighted). Both C57BL/6J and MRL/MpJ activated the same top 5 pathways with positive z-scores, including actin cytoskeleton signaling and integrin signaling pathways relative to respective uninjured groups (Fig. 5B). Interestingly, actin cytoskeleton signaling and integrin signaling pathways were also listed amongst the top 5 pathways in C57BL/6J injured tendons relative to MRL/MpJ injured tendons (Fig. 5C, z-score 2.294 and 2.985, respectively). This was also observed in C57BL/6J uninjured tendons relative to MRL/MpJ uninjured tendons (Fig. 5C).

**Fig. 5.**
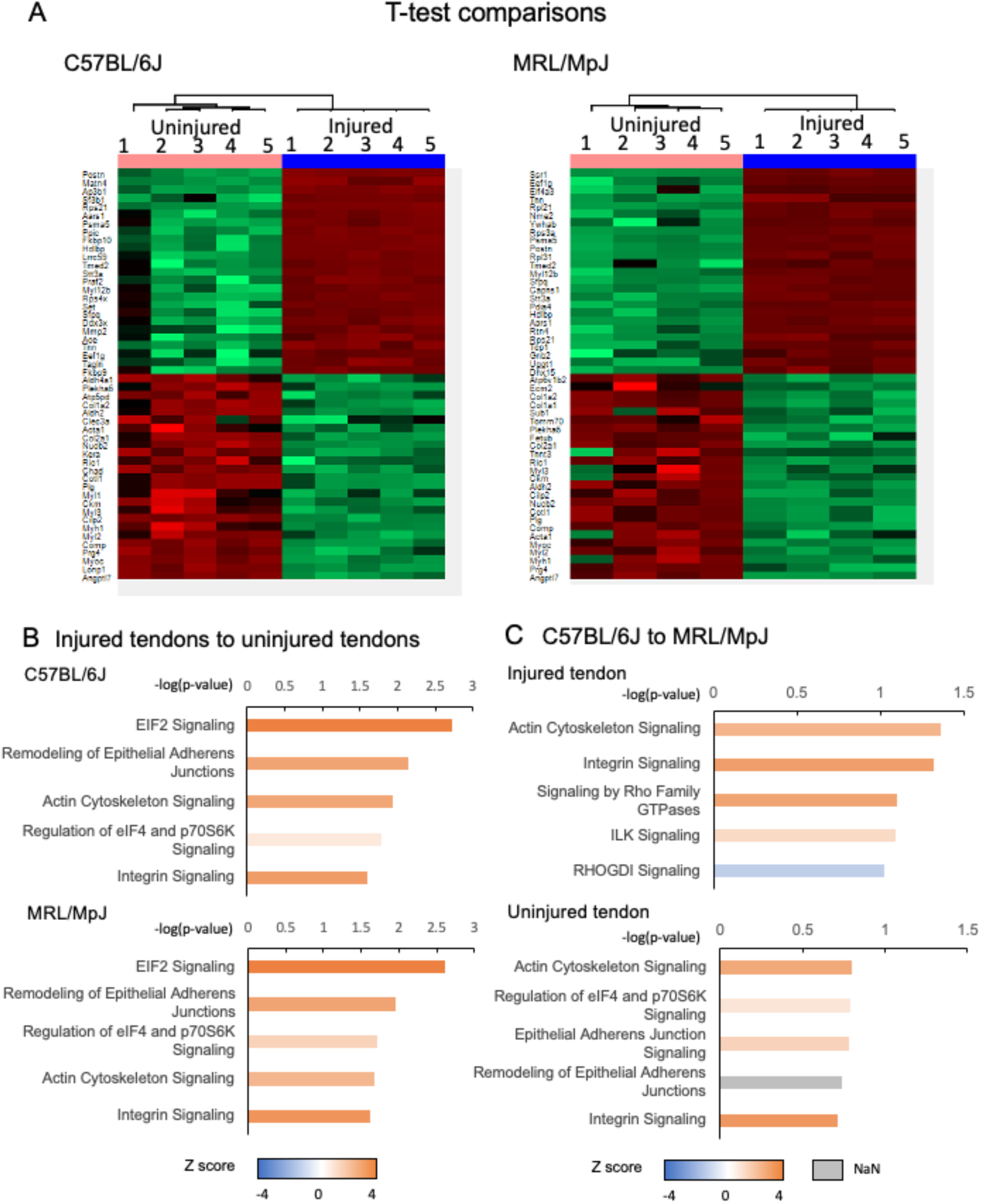
Proteomics analysis of injured tendons of C57BL/6J and MRL/MpJ mice. A complete transverse transection was made at the midpoint of the left Achilles tendon in 12-week-old male C57BL/6J (blue) and MRL/MpJ (red) mice. Four weeks after surgery, injured (left) and contralateral uninjured (right) tendons were harvested, and proteomics analysis was performed. (A) The top 50 upregulated and downregulated proteins in injured tendons samples in both C57BL/6J and MRL/MpJ male mice were well-clustered based on t-test comparisons. (B, C). The top 5 most altered signaling pathways were identified based on IPA analysis of differentially expressed proteins between injured and uninjured tendons of C57BL/6J and MRL/MpJ mice (B), and between C57BL/6J and MRL/MpJ tendons (C). Positive and negative z-scores represent upregulation and downregulation of the corresponding pathway, respectively.

## 4. Discussion

Our results indicate that tenotomized Achilles tendons of MRL/MpJ mice heal similarly as those of C57BL/6J mice, but with reduced heterotopic ossification than in C57BL/6J tendons by late stages of healing. Further, the injured tendons in MRL/MpJ male mice had lesser activation of actin cytoskeleton and integrin signaling pathways. These findings demonstrate MRL/MpJ mice possess limited capacity to regenerate injured tendons after complete rupture, but with reduced heterotopic ossification and activation of some molecular signaling pathways during tendon healing, such as cytoskeleton and integrin signaling, compared to C57BL/6J mice.

Superior healing by the MRL/MpJ mouse has been previously reported for patellar tendon injuries ^21^. After a 0.75 to 1 mm full-thickness focal defect in the patellar tendon, MRL/MpJ mice exhibit greater recovery of mechanical properties ^10^ and superior extracellular matrix realignment ^21^, compared to C57BL/6J mice. MRL/MpJ mice have also demonstrated improved recovery of mechanical strength after full-length and full-thickness central patellar tendon injuries ^11^, although pathology of their healing process has not been reported. In the Achilles tendon tenotomy model that we used here, C57BL/6J mice exhibit scar-mediated healing ^20^. Neo-tissue forms in the gap of the transected tendon in the first week after surgery, and the newly formed tissue gradually remodels over the next several weeks, with a decrease in cell density and increase in sagittal alignment of cells ^20^. However, the remodeled tissue still contains poorly aligned, thin collagen fibers ^8^ and undergoes chondroid degeneration and progressive heterotopic ossification 8 to 10 weeks after injury ^20,22^. Here, we found that after complete transection, MRL/MpJ injured Achilles tendons demonstrated fibrotic healing as observed in C57BL/6J mice, but the ectopic ossified mass was smaller in MRL/MpJ mice than in C57BL/6J mice. Furthermore, by 4 weeks of healing, there was no difference in the extent of recovery of mechanical properties between MRL/MpJ mice and C57Bl/6J mice regardless of sex. It appears that completely transected Achilles tendons in MRL/MpJ mice undergo scar-mediated healing, and do not have superior healing ability compared to C57BL/6J mice. However, additional investigations are needed, including quantitative analysis of histology, as well as mechanical testing and histological analysis at later stages.

In this study, we used a tendon injury model of complete transection of the Achilles tendon whereas previous studies have used a focal injury model of the patellar tendon that has shown regenerative healing ^10,21,23^. Considering the many differences between the tenotomized Achilles tendon injury model versus the patella tendon focal injury model, such as tendon type, injury size, and instability of injury site by motion, it is not possible to make generalized conclusions about the healing ability of MRL/MpJ tendons. In addition, the Achilles tendon injury model has limitations, including the development of ectopic bone formation, and the lack of maintaining the tendon ends in apposition following transection and during healing. Nevertheless, our results suggest that the regenerative capacity of tendons in the MRL/MpJ mouse may be limited depending on the type of injury as well as the anatomical location of the tendon. It has been suggested by others that MRL/MpJ regenerative healing extent depends on injury type or severity ^11^. Failure by the MRL/MpJ mouse to fully regenerate after injury has been demonstrated in other injured tissues, including the heart after ischemia-reperfusion injury ^9,24^, the skin after excisional injury ^25^, and articular cartilage after full-thickness injury ^26^. Here, we did not observe sex-specific differences in elastic modulus, ultimate stress, and ultimate strain of injured Achilles tendons 4 weeks after injury. This is consistent with previous findings that MRL/MpJ patellar tendon focal defects do not exhibit sex-specific differences in mechanical or material properties ^27^. However, male MRL/MpJ mice exhibit superior healing of articular cartilage full-thickness injuries compared to female MRL/MpJ mice ^26^. Interestingly, the heterotopic calcified volume in the injured Achilles tendons trended larger in males than in females, regardless of strain: 0.838+/- 0.600 mm^3^ versus 1.648 +/- 0.122 mm^3^ for C57BL/6J female versus male mice, respectively; and 0.209 +/-0.189 mm^3^ versus 0.588 +/- 0.348 mm^3^ for MRL/MpJ female versus male mice, respectively. In injured mouse Achilles tendons, heterotopic bone formation occurs via endochondral ossification at late healing stage ^20^. Taken together, male MRL/MpJ mice may have a stronger ability to generate cartilage tissue than female mice, resulting in the induction of larger volumes of cartilage-mediated ectopic bone in healing tendons.

To examine the molecular mechanisms involved in tendon healing, we performed proteomics analysis of injured tendons during the remodeling phase, 4 weeks after injury, because previous studies investigated the differences in the cells, matrix proteins, and signaling molecules between MRL/MpJ and C57BL/6J mouse patellar tendons during the inflammation and proliferation phases of early healing ^10–12,21,23^. Here, the altered signaling pathways in injured tendons during the remodeling phase were similar between MRL/MpJ and C57BL/6J mice, suggesting that in both strains, similar signaling pathways are altered upon injury, and similar molecular mechanisms are involved in scar formation and matrix remodeling. Intriguingly, we found that the MRL/MpJ male tendons showed less activation of the actin cytoskeleton and integrin signaling pathways than the C57BL/6J male tendons. These pathways govern cell adhesion, migration, proliferation, and matrix synthesis and organization ^28^. Upregulation of these pathways in response to injury is expected and is likely purposeful, especially during early healing phases after injury because inflammation and angiogenesis occur during the inflammatory phase, and reparative cells such as tendon progenitors and myofibroblasts migrate into and proliferate in the injured site during the proliferation phase ^29^. Actin cytoskeletal proteins and integrin adhesion complexes are essential molecules in regulation of migration, differentiation, matrix synthesis, and organization of myofibroblasts ^30^. It is of note that early upregulation of integrins and cellular adhesion genes have also been implicated in superior MRL/MpJ patellar tendon focal defect healing ^27^. Higher expression of *Itgb1* during the first 7 days post-injury was postulated to lead to improved cell and matrix alignment in MRL/MpJ healing patellar tendons, compared to C57BL/6J tendons. In contrast, during the remodeling phase, higher activity of these signaling pathways may be caused by or be associated with higher activity of fibrous tissue formation, leading to greater inhibition of structural recovery of collagen fibers and functional recovery of mechanical properties in C57BL/6J mice. Future studies should investigate the differences in the tendon healing processes between MRL/MpJ and C57BL/6J mice over a longer period in this tenotomy injury model. In particular, material property characterizations – including elastic modulus, ultimate strength, and collagen crosslink density – should be determined after additional longer healing durations to more fully evaluate long-term recovery of function and restoration of tissue quality. Quantitative assessments of histological characterizations, including fibrillar collagen density and alignment, cell and nuclear morphology, and integrin binding are needed to better inform the extent of scarless healing in MRL/MpJ mice. Identification of specific molecules that regulate the increased activity of the actin cytoskeleton and integrin signaling pathways and validation of their role in injured C57BL/6J tendon healing is also needed. Despite the unexpected discovery of limited MRL/MpJ tendon healing capacity observed in this study, these data suggest that lower activity of the actin cytoskeleton and integrin signaling pathways may be associated with a lower degree of heterotopic ossification. Complementing our current data with future proteomic characterizations of healing MRL/MpJ versus C57BL/6J female Achilles tendons, along with validation of specific protein targets, will provide more comprehensive insight into the sex-specific MRL/MpJ regenerative healing capacity for tendon injuries.

## Contributions

CKK and MEI designed the experiments. NT, KW, MW, HT, CDH, PR, and MEI conducted experiments; NT, DSS, JJ, NL, MAK, CKK, and MEI evaluated the results; and CKK and MEI prepared the manuscript. All authors have read and approved the final submitted manuscript.

## Acknowledgments

We thank D. Muthusamy and E. Mirdamadi for assistance with the DMA850.

## Supplementary Material

**Fig. S1.**
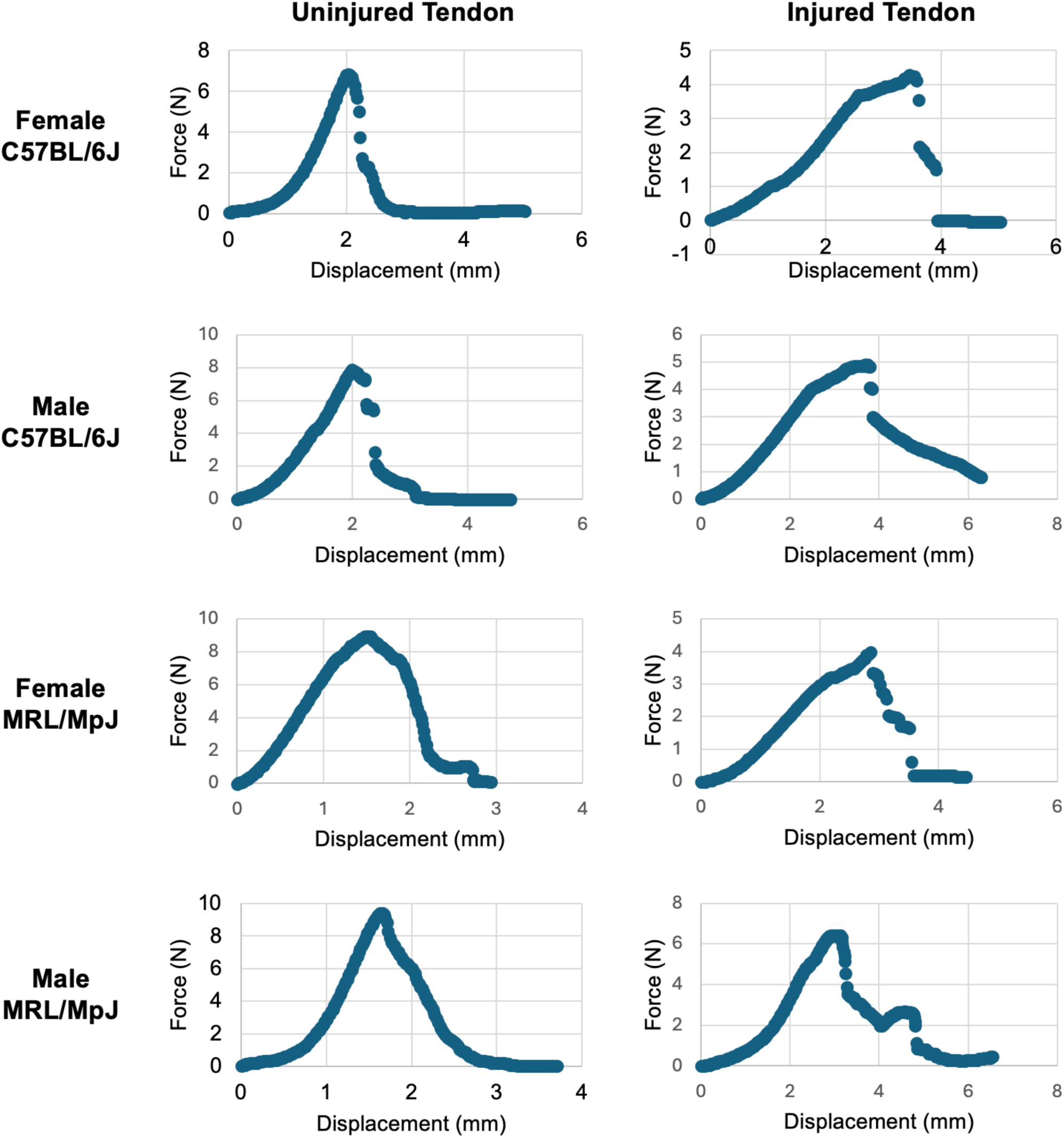
Representative force-displacement curves for uninjured and injured tendons of both strains and sexes.

**Fig. S2.**
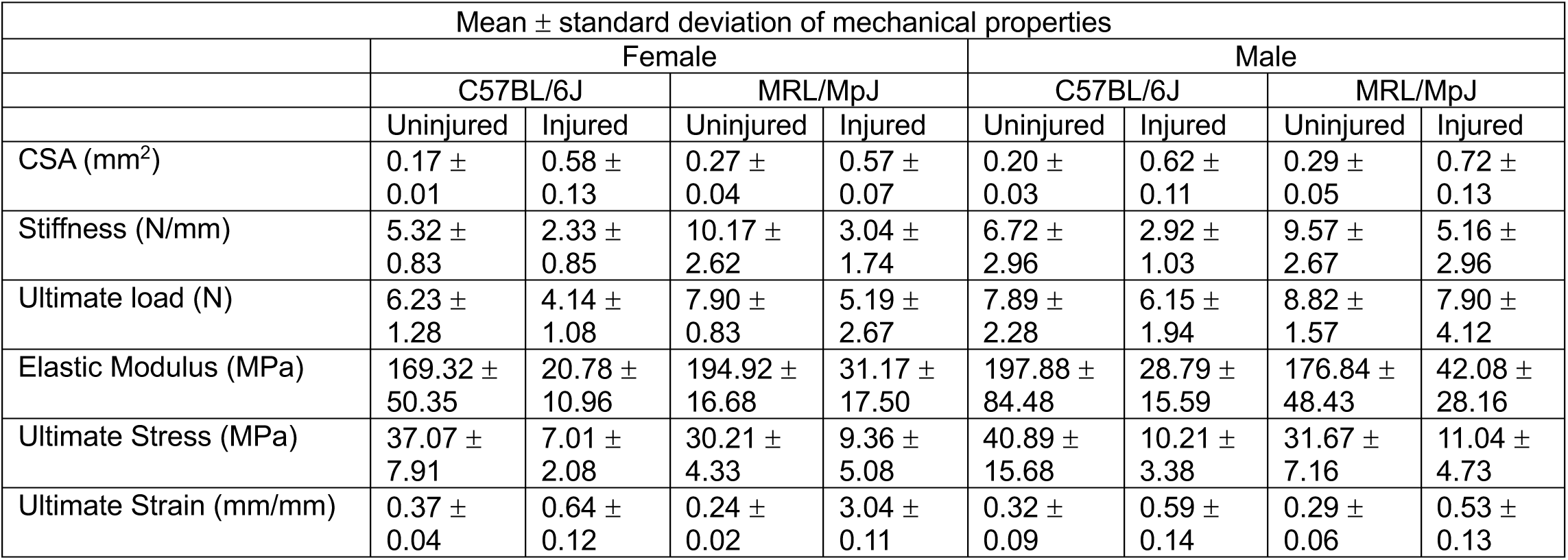
Averages and standard deviations of absolute values for cross-sectional area (CSA), stiffness, ultimate load, elastic modulus, ultimate stress, and ultimate strain of uninjured and injured tendons of C57BL/6J female, MRL/MpJ female, C57BL/6J male, and MRL/MpJ male mice.

